# Scalable Differentiable Folding for mRNA Design

**DOI:** 10.1101/2024.05.29.594436

**Authors:** Ryan Krueger, Max Ward

## Abstract

mRNA is an emerging therapeutic platform with applications ranging from vaccines to genome editing. However, there are an exponential number of mRNA sequences to deliver a given payload and the choice in nucleotide sequence largely determines stability and translation efficiency. There exist several computational approaches for optimizing mRNA sequences but these algorithms are limited in performance or the choice of optimization metric. In this work we describe a new mRNA design algorithm that overcomes both of these limitations and is based on differentiable folding, a recently developed paradigm for RNA design in which a probabilistic sequence representation is optimized via gradient-based methods. First, we present major improvements to the original differentiable folding algorithm that drastically reduce the memory overhead of the gradient calculation. Second, we formulate the mRNA design problem in the context of continuous sequences, requiring the generalization of existing metrics and careful treatment of constraints. Given this scaled algorithm and our mRNA design formalism, we then developed a generative deep learning approach that treats our differentiable folding algorithm as a module in a larger optimization pipeline to learn a network that samples optimized sequences. As a demonstration of our method, we optimize mRNA sequences via complex, therapeutically relevant objective functions.

## 1. Introduction

Ribonucleic acid (RNA) is a fundamental molecule in any biological organism and is therefore an attractive substrate for new therapeutic agents [1, 2, 3, 4]. Despite the advent of high-throughput sequencing and synthesis technology, experiments at the bench are costly and time-consuming. Therefore, computational modelling is used to obtain rapid approximations to experimentally determined structures [5, 6, 7, 8]. Accurate computational prediction of RNA structure from only the sequence is a widely studied problem [9, 10, 11, 12]. The inverse problem is equally important, but appears to be much harder. Instead of predicting the structural properties given the sequence, many biomedical applications require finding a sequence that has given structural properties. This is generally called an “inverse” or “backwards” problem. In the context of RNA, this problem is known as the “RNA design” problem.

There are two common kinds of computational RNA design problems: *structure design* and *mRNA design*. The structure design problem is to find a sequence that folds into a target structure under a computational model. The mRNA design problem is to find a sequence that maximizes vaccine suitability under a computational model, usually by considering stability and expression.

The structure design problem is well studied with algorithms proposed as early as 1994 [13], many new algorithms since then (see [14] for a summary), and humans solving gamified structure design puzzles [15]. Our focus in this work is on the mRNA design problem.

The mRNA design problem generally involves optimizing the protein coding region for a candidate vaccine sequence. Creating effective vaccine sequences involves many other considerations including the UTRs, capping, nucleoside modification, and addition of a poly(A) tail. However, since there is not yet a good computational objective for these, algorithms focus on optimizing the coding region while assuming templates are used for the other parts of the sequences mentioned previously. Many approaches optimize similarity to natural sequences [16, 17, 18, 19]. Recent approaches also consider the predicted stability of the sequence [20, 21, 22, 23]. Our major contribution is a new mRNA design algorithm that significantly advances the state of the art.

Our new algorithm builds upon Matthies *et al*.’s recently published differentiable RNA folding algorithm [24]. This algorithm computes the RNA partition function [25] but is differentiable, meaning it is compatible with gradient based optimization including neural networks. Matthies *et al*. [24] demonstrated the algorithm for structure design and provide several directions for future work including: improving the algorithm so it can be used on longer sequences, adapting it to mRNA design, and incorporating it into a neural network pipeline. The work presented here achieves all three of these goals to create a novel and effective mRNA design algorithm.

While preparing this manuscript, a preprint by Dai *et al*. appeared that optimizes mRNA sequences via differentiable folding [26]. Our methods are similar but with several key differences. Notably, our method extends the differentiable folding algorithm of Matthies *et al*. to build a neural network pipeline whereas Dai *et al*. do not. A full discussion of the similarities and differences are given in Section 3.1.

### 1.1 The mRNA Design Problem

There are many objectives one may want to optimize for mRNA design. For our purposes we use a simple but widely accepted formalization. This simplifies the problem to optimizing two objectives simultaneously, the stability of the sequence and the Codon Adaptation Index (CAI) [16]. Let us call this stability-CAI optimization. We give a generic formulation of stability-CAI optimization:

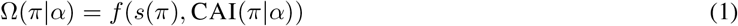

Ω(*π*|*α*) is the objective function where *α* is an amino acid sequence, *π* is a potential mRNA sequence. The function *s*(*π*) measures the stability of the sequence and CAI(*π*|*α*) is the CAI of the RNA sequence given amino acid sequence. The function *f* is used to combine the stability and the CAI. Different formulations have been given for *s* and *f*. CAI is a geometric mean:

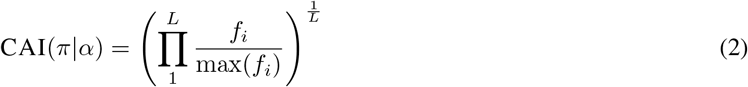

It is assumed that *π* corresponds to a valid codon sequence for *α*, so let 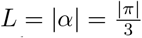. A valid codon sequence contains the right number of codons and each codon is a match for the corresponding amino acid. Let CAI(*π*|*α*) = 0 if *π* is not a valid codon sequence for *α*. Given a reference set of genes, the frequency of the *i*th codon encoded in *π* is denoted by *f*_*i*_. Correspondingly, max(*f*_*i*_) denotes the maximum frequency over all possible codons for the *i*th amino acid in *α*.

Terai, Kamegai, and Asai appear to be the first to propose stability-CAI optimization [23]. They proposed to minimize Ω(*π*|*α*) = MFE(*π*) × CAI(*π*|*α*)^*λ*^ where MFE(*π*) denotes the *minimum free energy* (MFE) for a sequence *π*, which is the free energy of the most stable structure in the ensemble [9]. Note that *λ* is an arbitrary weighting factor. Terai, Kamegai, and Asai proposed an algorithm to find a sequence with minimum MFE, but it could not simultaneously optimize CAI [23]. Later, Zhang *et al*. solved the problem [20]. They proposed LinearDesign, which minimizes Ω(*π*|*α*) = MFE(*π*) *− λ* log(CAI(*π*|*α*)). LinearDesign represented a breakthrough, as it was the first algorithm to effectively do stability-CAI optimization.

Other stability-CAI objectives have been proposed. Wayment-Steele *et al*. proposed *average unpaired probability* (AUP), an improved measure of stability, and Ribotree an algorithm for optimizing mRNA using a modified Monte Carlo tree search. AUP was experimentally determined to be a better measure of stability than MFE [22]. This is intuitive, since MFE only considers a single structure, but AUP considers the entire ensemble. Wayment-Steele *et* a*l*. also tried optimizing Δ*G*(ensemble), which is the free energy of the entire ensemble rather than just the most stable structure as in MFE. We call this *ensemble free energy* (EFE). EFE is a natural generalization of MFE, and is closely related to the stability measure we apply in our work. It should be observed that Wayment-Steele *et al*. reported a strong correlation between MFE and EFE, but only in sequences that were optimized for AUP alone. Our results show that the correlation breaks down with sufficient optimization, which means that EFE is likely to be an improvement over MFE in practice as well as in theory.

We chose to optimize the partition function *Z*_*π*_. A detailed explanation is given in Section 1.2. The succinct reasoning is that *Z*_*π*_ is closely related to EFE such that optimizing *Z*_*π*_ will also optimize EFE, but *Z*_*π*_ is more convenient to work with.

In the previously mentioned approaches a weighting factor *λ* is used to balance stability and CAI. Instead, we use set a threshold for CAI. That is, we optimize a sequence *π* to maximize stability subject to the condition that its CAI is at least *τ* .

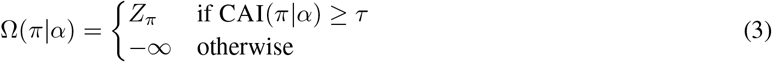

For our purposes, the mRNA design problem is to find an mRNA *π* that optimizes Equation (3) given an amino acid sequence *α*. The methods we present can likely be adapted to more complex objectives. These might include other measures of stability, such as AUP. They might also include other considerations relevant to a vaccine designer, such as 5’-leader region optimization [27], or uridine depletion [28]. However, Equation (3) is used here as it is at least as powerful as existing widely-used objectives and is simple enough for a concise presentation.

### 1.2 Ensemble Free Energy and the Partition Function

McCaskill’s famous algorithm [25] computes the RNA partition function. The partition function of a sequence *π* is defined as:

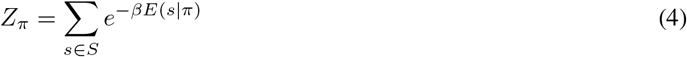

*S* is the set of all possible structures, *β* is a thermodynamic scaling factor, and *E*(*s*|*π*) is the free energy of the structure *s* with respect to the sequence *π*. EFE is related to the partition function:

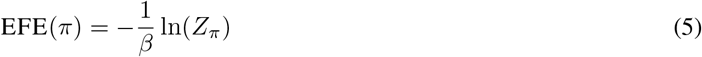

This means that we can maximize *Z*_*π*_ instead of EFE(*π*) for stability. For example, we could substitute EFE into Equation (3) and the optimization problem would not change.

We use the differentiable partition function calculation due to Matthies *et al*. [24]. It computes a generalised form of *Z*_*π*_, which allows us to optimize for sequences with high *Z*_*π*_.

### 1.3 Differentiable Partition Function

Matthies *et al*. introduce the notion of a differentiable partition function [24] and give a polynomial time algorithm to calculate it. This partition function differs from Equation (4) in two important ways. First, it is defined for a distribution of sequences Ψ.

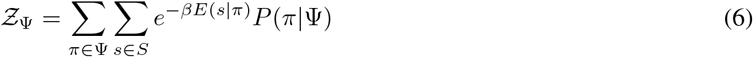

We use *P*(*π*|Ψ) to denote the probability of a sequence sampled from the distribution. In short, *Ƶ*_Ψ_ is an expected partition function: *Ƶ*_Ψ_ = 𝔼_*π∼*Ψ_[*Z*_*π*_].

It should be observed that the algorithm presented in [24] only handles Ψ constructed from a set of independent categorical distributions for each nucleotide. For a sequence of length *n*,

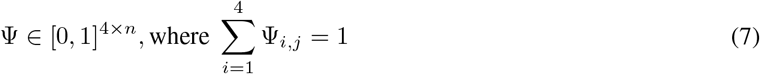

Let *π*_*j*_ ∈ [0, 4) be an integer denoting the nucleotide identity of the *j*th nucleotide in the sequence *π*.

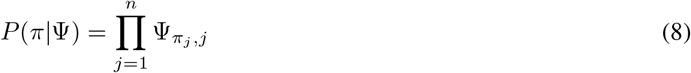

The second important difference between the work of Matthies *et al*. [24] and McCaskill’s partition function [25] is that the algorithm to compute *Ƶ*_Ψ_ is differentiable. This means that gradients can be taken with respect to the input, Ψ. So, we can define *Ƶ*_Ψ_ as the objective for maximization and optimize Ψ via gradient descent. The upshot of the optimization is a distribution Ψ with a high 𝔼_*π*∼Ψ_[*Z*_*π*_]; sampling from the resulting Ψ gives sequences with high partition functions. This is why [24] suggests mRNA design as a future direction for their algorithm.

There are two issues with the above approach. First, it only considers stability and not CAI. It optimizes the partition function, but it does not optimize CAI at all. Second, the algorithm of Matthies *et al*. ran out of memory for sequences whose length was greater than 50nt on an 80GB GPU [24]. We present new loss terms that allows to optimize a full stability-CAI objective. We also significantly improve the differentiable folding algorithm to increase the maximum length it can handle.

Matthies *et al*. speculated that the differentiable partition function algorithm could be incorporated into a deep learning pipeline, since gradients can be computed end-to-end [24]. Our mRNA design algorithm is the first to realize this approach. Instead of optimizing Ψ directly by gradient descent, we optimize a neural network that outputs Ψ. We find that this method significantly improves performance.

### 1.4 Our Contributions

The key contributions of this work are

1. We incorporate CAI into differentiable partition function optimization for mRNA via new loss functions. This enables the use of a stability-CAI objective.
2. We drastically improve the time and space performance of the differentiable partition function algorithm of [24]. Our improved algorithm is able to compute gradients for sequences with lengths up to 1250nt rapidly on a single GPU.
3. We incorporate our new and improved differentiable partition function algorithm into a deep learning pipeline. This allows more effective mRNA optimization.

## 2 Results

In this section, we demonstrate how to optimize mRNA sequences via differentiable folding and place our method in the context of existing design algorithms. First, we briefly describe a suite of improvements to scale the algorithm of Matthies *et al*. [24] to longer sequences. Second, we formulate the mRNA design problem at the level of continuous nucleotide sequences such that coding sequences may be optimized via differentiable folding. Thirdly, we introduce a generative deep learning approach that enables the efficient navigation of this complex optimization landscape. Lastly, we test our algorithm on several proteins by improving the results found by LinearDesign [20].

### 2.1 Algorithmic Improvements

The original implementation of differentiable folding was limited to sequences of up to 50 nucleotides in length [24]. The primary bottleneck for this algorithm was memory usage and GPUs have limited memory. To scale differentiable folding to experimentally-relevant mRNA lengths improvements to the algorithm were necessary. Note that the original McCaskill’s algorithm [25] required *O*(*n*^2^) memory for a sequence of *n* nucleotides, which is relatively frugal. However, continuous and differentiable folding first adds a large constant factor to deal with the continuous sequence, and also uses *O*(*n*^3^) memory since intermediate values are stored for backpropagation [24]. We developed a suite of algorithmic optimizations that taken together permit efficient gradient calculations for sequences of up to 1250 nucleotides on a single GPU.

The optimizations can be roughly split into three categories: better recursions, checkpointing, and numerical issues. The *better recursions* category includes improving the dynamic programming recursions used to compute the partition function. These all lead to constant factor improvements to speed and memory usage, although this constant factor was sometimes large. The treatment of coaxial stacks, terminal mismatches, dangling ends, and multiloop branches was changed, which saved a factor of 256. In addition, internal loops were also optimized saving a factor of 256.

We used *checkpointing* to exploit the time-memory tradeoff inherent to backpropagation. Gradient calculation by backpropagation entails saving intermediate values so that gradients can be calculated. Instead of storing all intermediate values, one can recompute them from checkpoints. We found that a judicious use of this could reduce our theoretical memory usage asymptotically from *O*(*n*^3^) to *O*(*n*^2.5^). The practical impact of this strategy is shown in Figure 2.

Improving *numerical issues* was the final step. With the algorithm able to handle longer lengths, numerical issues arose in the calculation. GPUs can generally only handle up to 64 bits of floating point precision. The dynamic programming algorithm takes a weighted sum of over the partition functions of all 4^*n*^ sequences and each partition function is a sum of an exponential number of structures [29]. This leads to very large values that can overflow. There are two methods generally used to mitigate overflow in RNA folding packages: (i) computing values in log-space (used in RNAstructure [6]) or (ii) per-nucleotide scaling (used in ViennaRNA [5]). We found that log-space is problematic for gradient calculations, because addition in log-space must be approximated, but the gradient of the approximation is not necessarily an approximation of the gradient. To avoid this issue, we chose to use per-nucleotide scaling.

### 2.2 mRNA Design as a Continuous Optimization Problem

We defined the mRNA design problem as finding the nucleotide coding sequence that maximizes Equation (3). Intuitively, Equation (3) maximizes our chosen measure of stability (i.e. the partition function) subject to a constraint that requires a minimum CAI. Since we address the mRNA design problem via continuous optimization, we must generalize these quantities to continuous sequences.

For a discrete RNA sequence *π*, CAI captures the codon usage bias for a given organism (see Section 1.1). We define the *expected CAI* for a distribution of sequences Ψ,

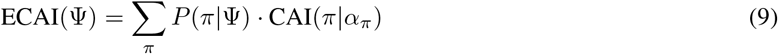

where CAI(*π*|*α*_*π*_) is defined as in Equation (2) and *α*_*π*_ is the amino acid sequence corresponding to *π*.

Note that we do not restrict the calculation of ECAI(*π*) to the target protein sequence *α* as this would underapproximate the CAI of the distribution of sequences. Instead, we compute an expectation over all codon sequences regardless of if they are valid for *α*. Importantly, ECAI(Ψ) can be computed efficiently by computing the contribution of each probabilistic codon independently.

The constraint that the nucleotide sequence must code for the target protein imposes an additional challenge for differentiable folding as the dynamic programming recursions are defined at the nucleotide level, rather than at the codon level. To control for this, we define an additional term describing the probability that a sequence sampled from Ψ codes for the target protein *α*:

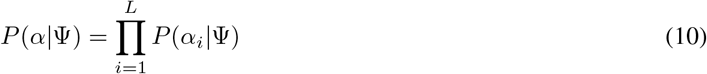

*P*(*α*_*i*_|Ψ) is the probability that Ψ codes for codon *α*_*i*_ and is defined as

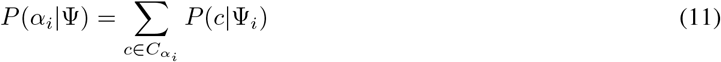

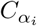 is the set of nucleotide triplets coding for codon *α*_*i*_, Ψ_*i*_ is the subsequence of Ψ corresponding to the *i*th codon, and *P*(*c*|Ψ_*i*_) is the probability of this subsequence coding for triplet *c*. Like ECAI, *P*(*α*|Ψ) can be computed efficiently by computing the probability of each codon subsequence independently.

We can then combine these two quantities to formulate a loss function for mRNA design over a distribution of sequences. Given an arbitrary metric of mRNA stability *O* where higher numbers represent greater stability, we represent the loss function of a distribution of sequences Ψ as

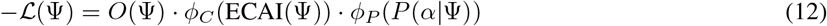

where *ϕ*_*C*_ and *ϕ*_*P*_ are activation functions that enforce minimum values of ECAI(Ψ) and *P*(*α*|Ψ), respectively. Both *ϕ*_*C*_ and *ϕ*_*P*_ “activate” if their values are below a threshold, which increases the loss. There are many choices of activation function; in our experiment we used a Leaky ReLU and a quadratic spline, see Section 4.1 for details.

We use take the product of loss terms instead of the sum to account for the different scales of each term. This is analogous to using a geometric mean instead of an arithmetic mean. The negative sign indicates the formulation of the optimization problem as a minimization problem.

In our experiments we use *O*(Ψ) = *Ƶ*_Ψ_ = ∑_*π*_ *p*(*π*|Ψ) · *Z*_*π*_.

Since we can compute *O*(Ψ) = *Ƶ*_Ψ_ using differentiable folding, Equation (12) is continuous and differentiable and we can directly optimize Ψ via gradient descent. Given an optimized Ψ, we obtain a final optimized nucleotide sequence by sampling discrete sequences from Ψ and choosing the sampled sequence that maximizes stability while satisfying the minimum value of CAI and coding for *α* as per our initial objective Equation (3) (see Figure 1).

**Figure 1:**
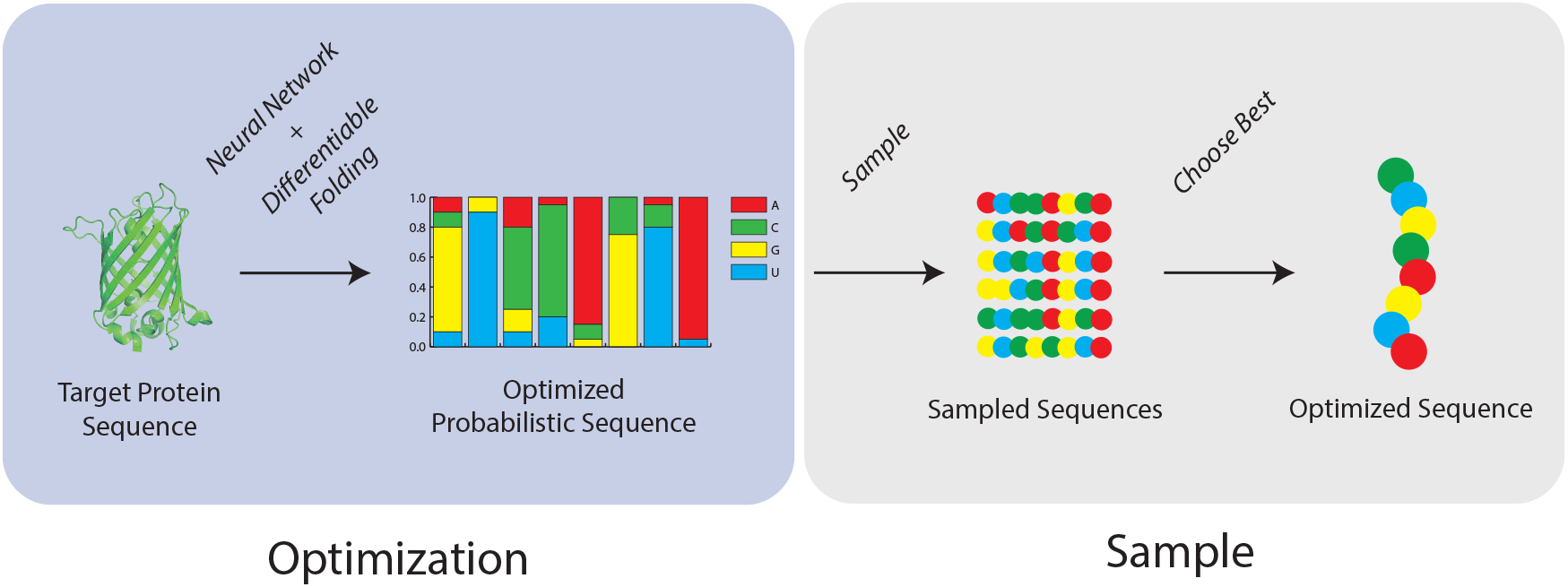
An overview of our method for mRNA design. Given a target protein sequence, we treat our differentiable folding algorithm as a module in a larger deep learning pipeline to optimize a probabilistic nucleotide sequence. We then sample final discrete sequences from the optimized distribution.

**Figure 2:**
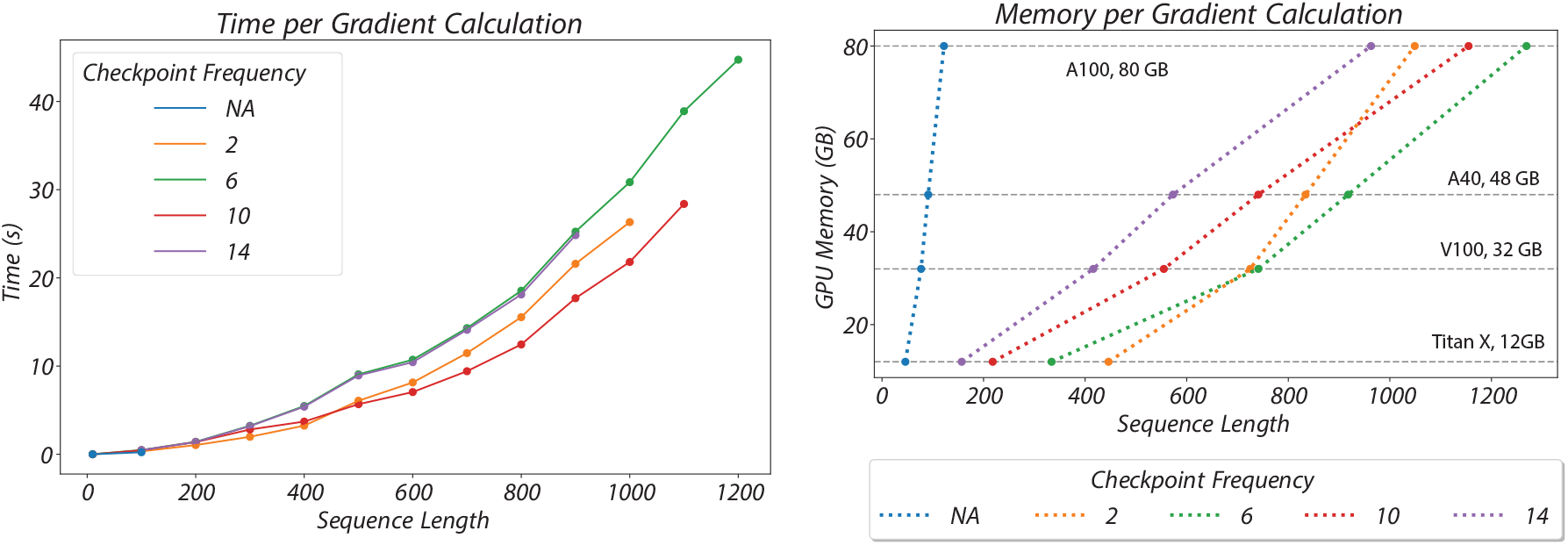
The time and memory cost with respect to sequence length for our optimized differentiable folding algorithm. On the left, we plot the time per gradient evaluation as a function of sequence length for different checkpointing frequencies. On the right, we plot the memory cost per gradient evaluation as a function of sequence lengths for the same set of checkpointing frequencies.

### 2.3 Generative Deep Learning Method

In its basic form, differentiable folding directly optimizes a continuous RNA sequence via gradient descent. However, as demonstrated in [24], vanilla gradient descent does not always find optimal solutions for structural RNA design. In the context of mRNA design, these problems will only be exacerbated by the non-convexities introduced by the loss terms for ECAI and the probability of sampling a valid sequence.

One popular solution to better navigating non-convex landscapes is overparameterization [30, 31]. Rather than optimizing over raw parameters, one can optimize over the parameters of a neural network whose outputs are the parameters that one wishes to optimize. Meanwhile, a defining advantage of differentiable folding is that it is gradient-based and therefore can be used as a module in a larger deep learning pipeline.

We devised a simple generative deep learning approach to improve upon the vanilla gradient descent performed in [24]. For a given RNA design problem, rather than optimizing over the continuous sequence itself, we optimize over the parameters of a fully connected neural network that outputs candidate continuous sequences as input to our folding algorithm. The input to the network is a random seed. See Figure 3 for an overview. Crucially, observe that such end-to-end training is only possible because the partition function calculation is itself differentiable; otherwise, we would need a large training set to learn from, which does not readily exist for RNA design.

**Figure 3:**
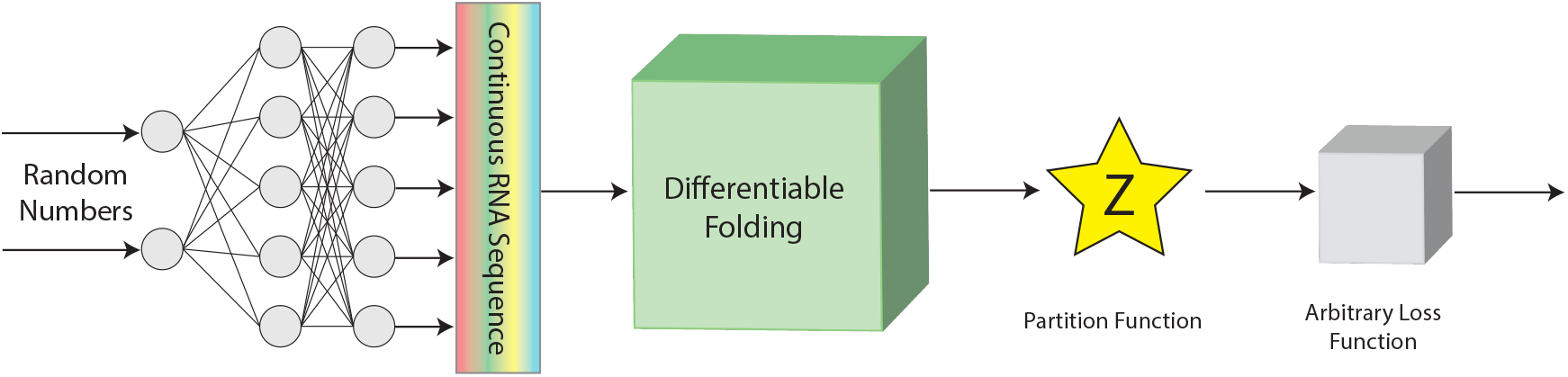
An overview of our method to train a generative model that produces probabilistic sequences that minimize a continuous loss function based on the partition function. We overparameterize the optimization problem by training a generative network to produce a sequence distribution that minimize the objective function. Once trained, discrete sequences can be sampled from the predicted sequence distribution.

**Figure 4:**
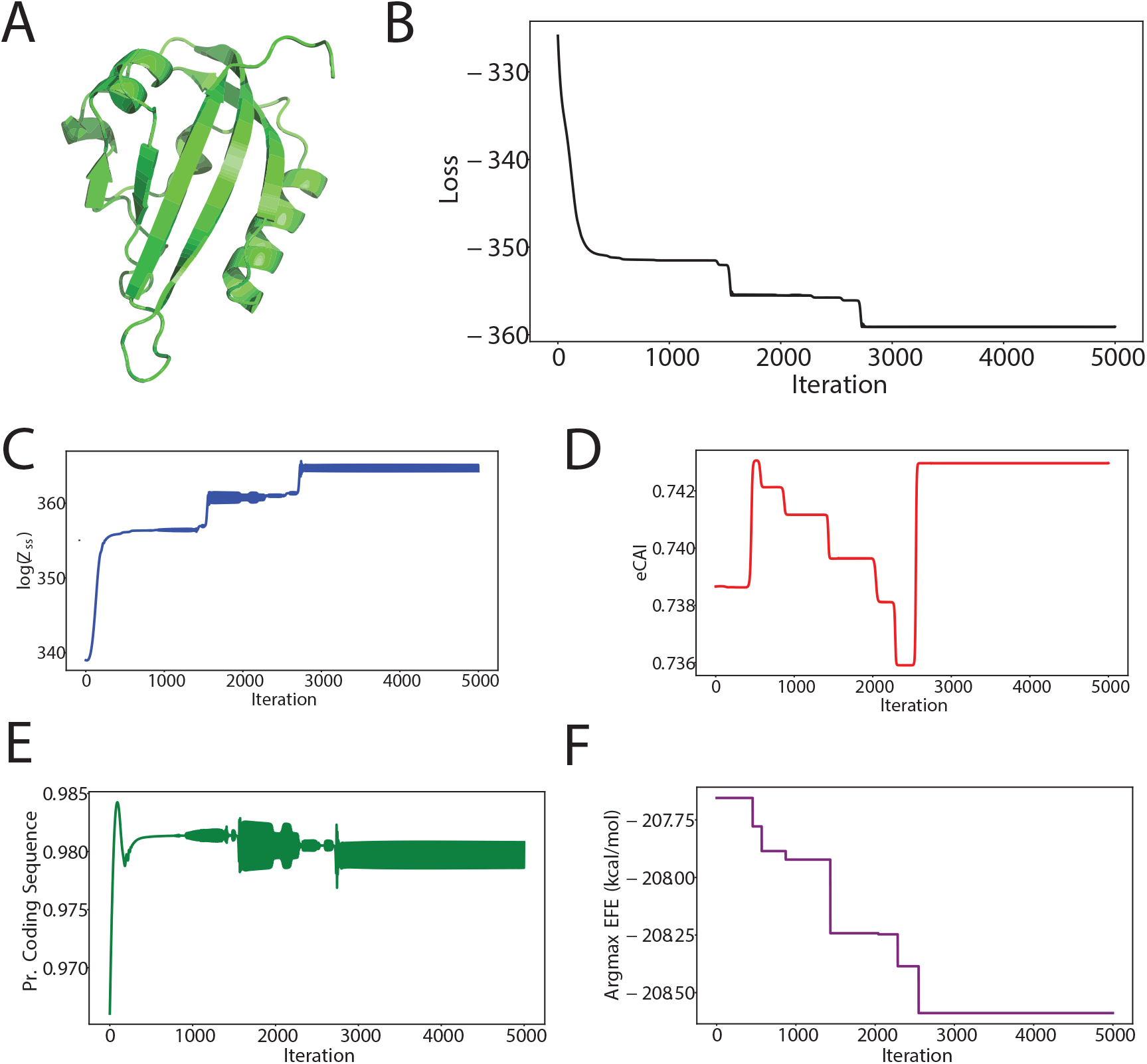
An example optimization of Mini GFP via our method with no CAI constraint and with *P*(*α*|Ψ) ≥ 0.95. (A) The predicted structure of Mini GFP via ColabFold [32]. The (B) loss, (C) log of the expected partition function *Ƶ*_*π*_, (D) expected CAI, (E) probability of sampling a valid coding sequence are plotted over time, and (F) argmax sequence EFE are plotted over time.

### 2.4 Example Optimizations

As a demonstration of our method, we optimized the stability of five benchmark protein sequences – four sequences from [21] as well as Mini-GFP. For each protein sequence, we considered two optimization problems: (i) unconstrained CAI, and (ii) CAI ≥ 0.8. We seeded our algorithm with a continuous embedding of the LinearDesign solution [20]. Our algorithm consistently improved the sequence, yielding values of ΔΔ*G* up to *−*1.11 kcal/mol. For each sequence, we only ran our method once. It should be observed that multiple runs can lead to better results. See Table 1 for these results and Section 4.1 for complete implementation details of these optimizations.

**Table 1:**
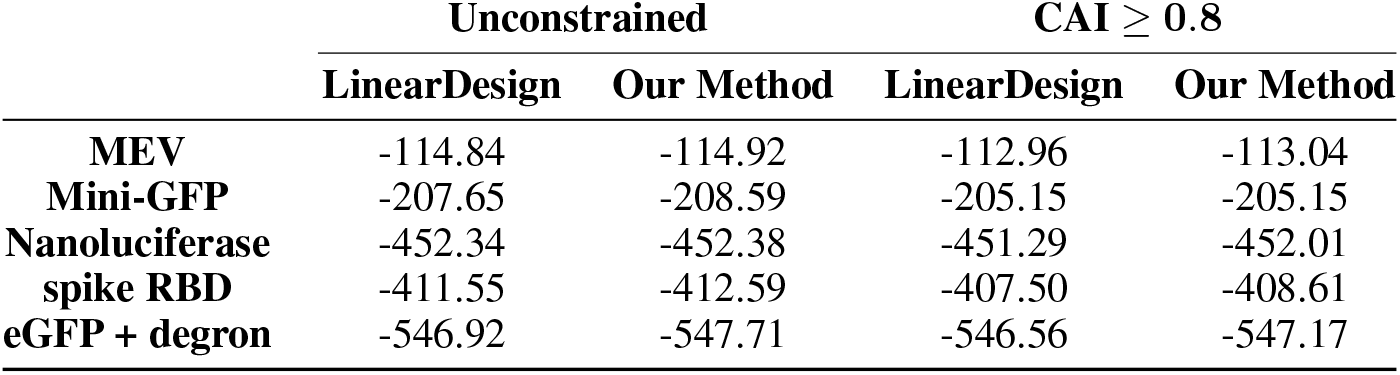
Optimized ensemble free energies of five benchmark mRNA sequences via our method and LinearDesign [20]. For each sequence, we considered two optimization targets: stability optimization (i) with unrestricted CAI and (ii) with CAI ≥ 0.8. The LinearDesign sequence was used as input to our method for each optimization run. All values are reported in kcal/mol.

One strength of our method highlighted by these example optimizations is its ability to consistently improve sequences in a single restart. This can be compared to the method of Dai *et al*. which requires a batch of random initializations for each sequence and can yield worse values of Δ*G* than the initial sequence whereas our method only requires a single initialization and will never make Δ*G* worse. One notable optimization is eGFP with CAI ≥ 0.8, for which our method found a sequence with lower EFE than the LinearDesign solution with unrestricted CAI.

## 3 Discussion

In this work, we present a new algorithm for mRNA design via differentiable folding. First, to make mRNA design tractable within the differentiable folding framework, we optimized the algorithm in [24] to achieve a ×25 increase in sequence length on a single GPU. Second, we defined an optimization framework for mRNA design with continuous sequences, requiring a notion of expected CAI and an additional penalty to constrain the set of nucleotide sequences to valid coding sequences. Third, given its gradient-based nature, we treat differentiable folding as a module in a generative deep learning pipeline that overparameterizes the space of continuous sequences. Our method improves state of the art performance on five benchmark protein sequences.

This work highlights the promise of differentiable folding compared to classical search algorithms for mRNA design. First, differentiable folding is *flexible*. Unlike existing approaches that are limited in their optimization targets, our method permits the use of arbitrary (continuous and differentiable) objective functions. While we used the partition function as a metric for stability, there are many alternative metrics that one could optimize via our method. Second, differentiable folding is *modular*. Although we used a simple generative scheme to improve our search algorithm, differentiable folding could be used as a module in *any* deep learning pipeline. Importantly, no data is required as the folding algorithm itself serves as a differentiable oracle. Third, differentiable folding is *performant*. Our results on five benchmark sequences, each requiring only a single optimization run, highlight the ability of gradient-based optimization to navigate complex optimization landscapes.

There is significant opportunity for future work in scaling our algorithm to longer sequences. One simple solution would be to distribute our algorithm across multiple GPUs, although distributing a dynamic programming algorithm and its gradient calculation across devices is challenging. An alternative solution would be to use one of several simplified models of RNA thermodynamics that achieve comparable accuracy without the various details of the nearest-neighbor model that impose a high memory cost (e.g., internal loops). Lastly, LinearPartition [33] approximates the partition function using beam-pruning with an efficient algorithm. This offers an attractive solution to the memory overhead imposed by differentiating through the entire partition function calculation. However, it is possible that not including low-probability structures in the partition function calculation would degrade the quality of the gradient signal, and implementing this algorithm on a GPU is non-trivial. Dai *et al*. have already implemented the linearized partition function and found some prelimiary results [26]. Their findings are discussed further in Section 3.1.

There is also opportunity to improve our search algorithm beyond a simple overparameterisation. Rather than training the weights of the network from scratch for each target mRNA, such a network could be pretrained to produce an optimized initial continuous sequences for a target mRNA. In addition to improving our integration with deep learning to better navigate sequence space, we could also increase the convexity of this optimization landscape by defining the recursions at the level of codons rather than nucleotides. Under our current formulation, for a mRNA sequence for *n* codons, the partition function sums over all 4^3*n*^ nucleotide sequences of which a maximum of only 6^*n*^ << 4^3*n*^ are valid coding sequence (as an amino acid corresponds to at most 6 codons). Restricting the calculation of the partition function to only valid coding sequences would significantly increase the convexity of sequence space. However, this is likely to add a large constant factor to the cost of computing the differentiable partition function and keeping track of CAI [20, 34], particularly if implemented on a GPU.

### 3.1 Comparison to Dai *et al*. [26]

During the preparation of this manuscript a preprint by Dai *et al*. was published that optimizes mRNA sequences similarly, i.e. via differentiable folding [26]. There is overlap between our work, but also some major differences. Both methods optimize the stability of mRNA sequences measured by expected partition function using a differentiable folding algorithm. There are several differences.

First, our method builds upon the differentiable folding algorithm of Matthies *et al*. [24]. Dai *et al*. do not use the same algorithm but instead implement a new folding algorithm based on LinearPartition [33] and LinearDesign [20]. The upshot of using different folding algorithms is that Dai *et al*. optimize an approximate partition function, whereas we optimize the complete partition function. This makes their method use fewer resources, but at the cost of lower quality gradients. In addition, our algorithm runs on GPU but the Dai *et al*. method is CPU-only. These folding algorithms are conceptually different as well as practically different. We operate at the level of nucleotides and use Equation (11) to enforce valid codon sequences, but Dai *et al*. operate at the level of codons.

Second, Dai *et al*. do not incorporate differentiable folding as a module in a larger deep learning architecture. In contrast, one of our major contributions is building a deep learning pipeline to optimize sequences. To achieve good results, Dai *et al*. used multiple initializations, whereas we only used a single run to generate our results.

Finally, we optimize a stability-CAI objective using the expected CAI loss term Equation (9). Dai *et al*. only optimize stability.

### 3.2 Conclusion

Differentiable folding presents a promising avenue towards next-generation mRNA design—not only can it efficiently navigate sequence space for arbitrary objective functions, but it can be combined with state of the art deep learning methods for superior performance. The high memory cost of differentiation presents the largest barrier for various practical applications, but these limitations can likely be overcome with standard optimizations. While not explored in this work, differentiable folding is amenable to complex mRNA design strategies, such as incorporating UTRs and imposing structure constraints on subsequences. This flexibility presents great opportunity for not only design with respect to existing metrics, but also the development of new metrics for mRNA stability and therefore a deeper understanding of mRNA thermodynamics.

## 4 Methods

### 4.1 Sequence Optimization

In Table 1, we report the optimized ensemble free energies via LinearDesign [20] and our method for five benchmark protein sequences. To generate results via LinearDesign for unconstrained CAI, we ran the publicly available executable with default settings and *λ* = 0. To find the optimal coding sequence subject to CAI ≥ 0.8 with LinearDesign, we performed binary search over *λ* to find the minimal *λ* (and therefore the minimum MFE) corresponding to a sequence satisfying the CAI constraint.

For the loss terms *ϕ*_*C*_ and *ϕ*_*P*_, we use two different activation functions and thresholds. For *ϕ*_*C*_, we use a Leaky ReLU with with a slope of 1.0 for ECAI < 0.8 and a slope of 0.05 for ECAI ≥ 0.8. For *ϕ*_*P*_, we use a quadratic spline with a quadratic coefficient of 50 for *P*(*α*|Ψ) < 0.95 and a linear slope of 0.025 for *P*(*α*|Ψ) ≥ 0.95.

All optimizations via our method were performed in JAX [35] on an 80 GB NVIDIA A100. To ensure that the continuous sequence is properly normalised, in practice we optimize a set of logits that are normalised via a softmax operator at each optimization step. Each optimization is performed in three steps:

1. Initialization of logits
2. Neural network pretraining
3. Sequence optimization

Logits are initialisized via the LinearDesign solution. For a nucleotide sequence of length *n*, the logits are initialised as 10 × onehot(seq_*LD*_) + 10 where seq_*LD*_ is the LinearDesign solution and onehot(seq_*LD*_) is the *n* × 4 one-hotted array corresponding to this discrete sequence.

Next, we pretrain a fully-connected neural network to predict this initial logits given a fixed random seed. We use a network with 6 layers, each consisting of 4000 features, and Leaky ReLU as an activation function. We define the pretraining loss function as the MSD between the predicted logits and the target initial logits. Pretraining is performed via 250 iterations with an Adam optimizer and a learning rate of 0.0001.

Finally, we apply gradient descent to the parameters of this pretrained network to predict logits that minimise our target objective function for mRNA. Each optimization was run for 5000 iterations (or until convergence) with a LAMB optimizer and a learning rate of 10^*−*5^.

Note that we tried to match the thermodynamic model used by LinearDesign [20]. Since their method is closed source, it is hard to be certain. However, they appear to use the same model as ViennaRNA [5] with the -d0 flag enabled, which does not count dangling ends.

## Acknowledgments

We thank Michael Brenner for his mentorship and support. This material is based upon work supported by the National Science Foundation under Grant No. UWSC13223.

## A mRNA Sequences

In this section we provide the amino acid and nucleotide sequences corresponding to Table 1. For each protein, in addition to providing the amino acid sequence, we provide the LinearDesign nucleotide sequence and the nucleotide sequence optimized via our method.

### Multi-epitope Vaccine

#### Amino Acid Sequence

~~~
MGGSGGSGYQPYRVVVLGGSGGSPYRVVVLSFGGSGGSLSPRWYFYY
~~~

#### Linear Design (unrestricted). CAI: 0.715

~~~
AUGGGUGGCAGCGGGGGCUCAGGCUACCAGCCCUACCGAGUGGUAGUACUCGGCGGGUCGGGCGGGAGCCCGUAC
CGCGUCGUAGUGCUAUCAUUCGGUGGGAGCGGUGGUAGCCUGAGCCCCCGCUGGUACUUCUACUAC
~~~

#### Our Method (unrestricted). CAI: 0.708

~~~
AUGGGUGGCAGCGGGGGCUCAGGCUACCAGCCCUACCGAGUGGUAGUACUCGGCGGGUCGGGCGGGAGCCCGUAC
CGCGUCGUAGUGCUAUCAUUCGGUGGGAGCGGUGGUAGCCUGAGCCCCCGCUGGUACUUCUAUUAU
~~~

#### Linear Design (CAI ≥ 0.8). CAI: 0.825

~~~
AUGGGUGGCAGCGGGGGCUCAGGCUACCAGCCCUACCGCGUCGUGGUUCUGGGCGGCAGCGGCGGUAGCCCCUAC
CGCGUUGUCGUCCUGAGCUUCGGCGGGUCGGGCGGUAGCCUGAGCCCCCGCUGGUACUUCUACUAC
~~~

#### Our Method (CAI ≥ 0.8). CAI: 0.821

~~~
AUGGGUGGCAGCGGGGGCUCAGGCUACCAGCCCUACCGCGUCGUGGUUCUGGGCGGCAGCGGCGGUAGCCCCUAC
CGCGUUGUCGUCCUGAGCUUCGGCGGGUCGGGCGGUAGCCUGAGCCCCCGCUGGUACUUCUAUUAC
~~~

### Mini GFP

#### Amino Acid Sequence

~~~
MEKSFVITDPWLPDYPIISASDGFLELTEYSREEIMGRNARFLQGPETDQATVQKIRDAIRDRRPTTVQLINYTK
SGKKFWNLLHLQPVFDGKGGLQYFIGVQLVGSDHV
~~~

#### Linear Design (unrestricted). CAI: 0.739

~~~
AUGGAGAAAUCCUUCGUGAUCACUGACCCCUGGCUGCCGGACUACCCGAUAAUCUCGGCUUCGGAUGGUUUCCUU
GAGCUGACCGAGUAUAGUCGGGAGGAGAUCAUGGGUCGCAACGCGCGUUUUCUGCAGGGGCCUGAGACGGAUCAG
GCCACCGUGCAGAAGAUCCGCGAUGCGAUCCGUGAUCGCCGCCCGACUACGGUUCAGCUCAUAAACUACACGAAG
UCGGGGAAGAAGUUCUGGAAUCUUCUUCACCUGCAGCCCGUCUUUGACGGCAAAGGCGGGCUGCAGUACUUUAUC
GGGGUCCAGCUAGUGGGCAGUGAUCACGUG
~~~

#### Our Method (unrestricted). CAI: 0.743

~~~
AUGGAGAAAUCCUUCGUGAUCACUGACCCCUGGCUGCCCGACUACCCGAUAAUCUCGGCUUCGGAUGGUUUUCUU
GAGCUGACCGAGUAUAGUCGGGAGGAGAUCAUGGGUCGCAACGCGCGUUUUCUGCAGGGGCCUGAGACGGAUCAG
GCCACCGUGCAGAAGAUCCGCGAUGCGAUCCGUGAUCGCCGCCCGACUACGGUUCAGCUCAUAAACUAUACGAAG
UCGGGGAAGAAGUUUUGGAAUCUUCUUCAUCUGCAGCCCGUCUUUGACGGCAAAGGCGGGCUGCAGUAUUUUAUC
GGGGUGCAGCUAGUGGGCAGUGAUCACGUG
~~~

#### Linear Design (CAI ≥ 0.8). CAI: 0.834

~~~
AUGGAGAAAUCCUUCGUGAUCACUGACCCCUGGCUGCCCGACUACCCCAUUAUAAGUGCCUCCGACGGCUUUCUU
GAGCUGACGGAGUAUAGUCGGGAGGAGAUCAUGGGUCGCAACGCGCGUUUUCUGCAGGGGCCUGAGACCGAUCAG
GCCACCGUGCAGAAGAUCCGCGAUGCGAUCCGUGAUCGCCGCCCGACUACGGUGCAGCUGAUUAACUACACCAAG
AGCGGCAAGAAGUUCUGGAAUCUGCUGCACCUGCAGCCCGUCUUUGACGGCAAAGGCGGGCUGCAGUACUUUAUU
GGGGUGCAGCUAGUGGGCAGUGAUCACGUG
~~~

#### Our Method (CAI ≥ 0.8). CAI: 0.834

~~~
AUGGAGAAAUCCUUCGUGAUCACUGACCCCUGGCUGCCCGACUACCCCAUUAUAAGUGCCUCCGACGGCUUUCUU
GAGCUGACGGAGUAUAGUCGGGAGGAGAUCAUGGGUCGCAACGCGCGUUUUCUGCAGGGGCCUGAGACCGAUCAG
GCCACCGUGCAGAAGAUCCGCGAUGCGAUCCGUGAUCGCCGCCCGACUACGGUGCAGCUGAUUAACUACACCAAG
AGCGGCAAGAAGUUCUGGAAUCUGCUGCACCUGCAGCCCGUCUUUGACGGCAAAGGCGGGCUGCAGUACUUUAUU
GGGGUGCAGCUAGUGGGCAGUGAUCACGUG
~~~

### JEV+Nluc

#### Amino Acid Sequence

~~~
MWLVSLAIVTACAGAMAVYPYDVPDYAGYPYDVPDYAGSYPYDVPDYAGSGVFTLEDFVGDWRQTAGYNLDQVLE
QGGVSSLFQNLGVSVTPIQRIVLSGENGLKIDIHVIIPYEGLSGDQMGQIEKIFKVVYPVDDHHFKVILHYGTLV
IDGVTPNMIDYFGRPYEGIAVFDGKKITVTGTLWNGNKIIDERLINPDGSLLFRVTINGVTGWRLCERILA
~~~

#### Linear Design (unrestricted). CAI: 0.791

~~~
AUGUGGCUAGUGUCGCUAGCCAUCGUGACCGCUUGCGCAGGCGCCAUGGCCGUGUACCCGUAUGACGUCCCCGAU
UAUGCAGGCUACCCGUACGAUGUGCCGGAUUACGCGGGGUCAUACCCCUAUGACGUUCCAGAUUAUGCCGGCUCU
GGCGUCUUCACCCUCGAAGACUUCGUGGGUGACUGGCGCCAGACGGCCGGCUAUAAUCUGGACCAGGUUCUGGAG
CAGGGAGGAGUGUCCUCCCUGUUCCAGAACCUGGGCGUCAGUGUGACCCCGAUCCAGCGCAUCGUGCUGUCCGGG
GAGAACGGCCUCAAAAUCGAUAUUCAUGUUAUCAUUCCAUACGAGGGUCUCAGCGGCGAUCAGAUGGGGCAGAUC
GAGAAGAUCUUCAAGGUGGUCUACCCCGUCGAUGACCAUCAUUUCAAAGUGAUCCUUCACUAUGGAACGUUGGUC
AUCGACGGGGUGACCCCUAACAUGAUAGAUUAUUUCGGUCGGCCCUAUGAGGGGAUCGCCGUGUUUGACGGUAAG
AAGAUUACCGUCACUGGGACCCUCUGGAAUGGUAACAAGAUUAUCGAUGAGAGGCUGAUCAACCCGGACGGUAGC
CUGCUUUUUCGGGUGACGAUCAACGGGGUCACGGGCUGGCGCCUGUGCGAGCGGAUCCUCGCG
~~~

#### Our Method (unrestricted). CAI: 0.782

~~~
AUGUGGCUAGUGUCGCUAGCCAUCGUGACCGCUUGCGCAGGCGCCAUGGCCGUGUACCCGUAUGACGUCCCCGAU
UAUGCAGGCUACCCGUACGAUGUGCCGGAUUAUGCGGGGUCAUACCCAUAUGACGUUCCAGAUUAUGCCGGCUCU
GGCGUCUUCACCCUCGAAGACUUCGUGGGUGACUGGCGCCAGACGGCCGGCUAUAAUCUGGACCAGGUUCUGGAG
CAGGGAGGAGUGUCCUCCCUGUUCCAGAACCUGGGCGUCAGUGUGACCCCGAUCCAGCGCAUCGUACUGUCCGGG
GAGAACGGCCUCAAAAUCGAUAUUCAUGUUAUCAUUCCAUACGAGGGUCUCAGCGGCGAUCAGAUGGGGCAGAUC
GAGAAGAUCUUCAAGGUGGUCUACCCCGUCGAUGACCAUCAUUUCAAAGUGAUCCUUCACUAUGGAACGUUGGUC
AUCGACGGGGUGACCCCUAACAUGAUAGAUUAUUUCGGUCGGCCCUAUGAGGGGAUCGCCGUUUUUGACGGUAAG
AAGAUUACCGUCACUGGGACCCUCUGGAAUGGUAACAAGAUUAUCGAUGAGAGGCUGAUCAACCCGGACGGUAGC
CUGCUUUUUCGGGUGACGAUCAACGGGGUCACGGGCUGGCGCCUGUGCGAGCGGAUCCUCGCG
~~~

#### Linear Design (CAI ≥ 0.8). CAI: 0.821

~~~
AUGUGGCUGGUGUCACUGGCCAUCGUAACCGCUUGCGCAGGCGCCAUGGCCGUGUACCCGUAUGACGUCCCCGAU
UAUGCAGGCUACCCGUACGAUGUGCCGGAUUACGCGGGGUCAUACCCCUAUGACGUUCCAGAUUAUGCCGGCUCU
GGCGUCUUCACCCUCGAAGACUUCGUGGGUGACUGGCGCCAGACCGCCGGCUAUAAUCUGGACCAGGUUCUGGAG
CAGGGAGGAGUGUCCUCCCUGUUCCAGAACCUGGGCGUCAGUGUGACCCCGAUCCAGCGCAUCGUGCUGUCCGGG
GAGAACGGCCUCAAAAUCGAUAUUCAUGUUAUCAUUCCAUACGAGGGCCUGUCCGGUGAUCAGAUGGGGCAGAUC
GAGAAGAUCUUCAAGGUGGUCUACCCCGUCGAUGACCAUCAUUUCAAGGUGAUCCUUCACUAUGGAACGUUGGUC
AUCGACGGGGUGACCCCUAACAUGAUAGAUUAUUUCGGUCGGCCCUAUGAGGGCAUUGCCGUCUUCGACGGCAAG
AAGAUCACCGUGACAGGCACUCUGUGGAAUGGUAACAAGAUUAUCGAUGAGAGGCUGAUCAACCCGGACGGUAGC
CUGCUUUUUCGGGUGACGAUCAACGGGGUCACGGGCUGGCGCCUGUGCGAGCGGAUACUGGCC
~~~

#### Our Method (CAI ≥ 0.8). CAI: 0.815

~~~
AUGUGGCUGGUGUCACUGGCCAUAGUAACCGCUUGCGCAGGCGCCAUGGCCGUGUACCCGUAUGACGUCCCCGAU
UAUGCAGGCUACCCGUACGAUGUGCCGGAUUACGCGGGGUCAUACCCCUAUGACGUUCCAGAUUAUGCCGGCUCU
GGCGUCUUCACCCUCGAAGACUUCGUGGGUGACUGGCGCCAGACAGCCGGCUAUAAUCUGGACCAGGUUCUGGAG
CAGGGAGGAGUGUCCUCCCUGUUCCAGAACCUGGGCGUCAGUGUGACCCCGAUCCAGCGCAUCGUGCUGUCCGGG
GAGAACGGCCUCAAAAUCGAUAUUCAUGUUAUCAUUCCAUAUGAGGGCCUGUCCGGUGAUCAGAUGGGGCAGAUC
GAGAAGAUCUUCAAGGUGGUCUACCCCGUCGAUGACCAUCAUUUCAAGGUGAUCCUUCACUAUGGAACGUUGGUC
AUCGACGGGGUGACCCCUAACAUGAUAGAUUAUUUCGGUCGGCCCUAUGAGGGCAUUGCCGUCUUCGACGGCAAG
AAGAUCACCGUGACAGGCACUCUGUGGAAUGGUAACAAGAUUAUCGAUGAGAGGCUGAUCAACCCGGACGGUAGC
CUGCUUUUUCGGGUGACGAUCAACGGGGUCACGGGCUGGCGCCUGUGCGAGCGGAUACUGGCC
~~~

### JEV+RBD

#### Amino Acid Sequence

~~~
MWLVSLAIVTACAGATNLCPFGEVFNATRFASVYAWNRKRISNCVADYSVLYNSASFSTFKCYGVSPTKLNDLCF
TNVYADSFVIRGDEVRQIAPGQTGKIADYNYKLPDDFTGCVIAWNSNNLDSKVGGNYNYLYRLFRKSNLKPFERD
ISTEIYQAGSTPCNGVEGFNCYFPLQSYGFQPTNGVGYQPYRVVVLSFELLHAPATVCGP
~~~

#### Linear Design (unrestricted). CAI: 0.759

~~~
AUGUGGCUCGUAAGCCUGGCCAUCGUGACGGCGUGUGCAGGAGCUACGAAUCUCUGCCCGUUUGGCGAGGUGUUC
AACGCCACGCGGUUCGCUAGUGUGUAUGCGUGGAAUCGCAAGCGCAUUAGCAACUGCGUGGCGGAUUAUUCAGUU
UUGUACAACUCAGCCUCGUUCAGCACGUUCAAGUGCUACGGGGUGAGUCCUACAAAGCUGAAUGAUCUCUGCUUC
ACCAACGUCUACGCGGAUAGCUUCGUUAUCCGCGGGGACGAGGUGAGGCAGAUAGCACCUGGCCAGACGGGCAAG
AUAGCCGAUUAUAAUUAUAAGUUGCCCGACGACUUUACUGGCUGCGUUAUUGCUUGGAAUAGCAAUAACUUAGAC
AGUAAAGUCGGGGGCAACUAUAAUUAUCUGUAUCGGCUGUUUCGGAAGAGCAAUUUGAAGCCCUUCGAGCGUGAU
AUAUCUACGGAGAUAUAUCAGGCUGGAUCCACCCCGUGCAACGGGGUGGAGGGCUUCAAUUGCUACUUCCCACUG
CAAUCCUAUGGGUUCCAGCCCACCAACGGGGUGGGCUACCAGCCCUAUAGGGUUGUAGUGCUGAGUUUCGAGCUC
CUGCACGCGCCGGCCACGGUUUGCGGGCCA
~~~

#### Our Method (unrestricted). CAI: 0.747

~~~
AUGUGGCUCGUAAGCCUGGCCAUCGUGACGGCGUGUGCAGGAGCUACGAAUCUCUGCCCGUUUGGCGAGGUGUUU
AACGCCACGCGGUUUGCUAGUGUGUAUGCGUGGAAUCGCAAGCGCAUUAGCAACUGCGUGGCGGAUUAUUCAGUU
UUGUACAACUCAGCCUCGUUUAGCACGUUCAAGUGCUACGGGGUGAGUCCUACAAAGCUGAAUGAUCUCUGCUUC
ACCAACGUCUACGCGGAUAGCUUCGUUAUCCGCGGAGACGAGGUGAGGCAGAUAGCACCUGGCCAGACGGGCAAG
AUAGCCGAUUAUAAUUAUAAGUUGCCCGACGACUUUACUGGCUGCGUUAUUGCUUGGAAUAGCAAUAACUUAGAC
AGUAAAGUCGGGGGCAACUAUAAUUAUUUGUAUCGGCUAUUUCGGAAGAGCAAUUUGAAGCCCUUCGAGCGUGAU
AUAUCUACGGAGAUAUAUCAGGCUGGAUCCACCCCGUGCAACGGGGUGGAGGGCUUCAAUUGCUACUUCCCACUG
CAAUCCUAUGGGUUUCAGCCCACCAACGGGGUGGGCUACCAGCCCUAUAGGGUUGUAGUGCUGAGUUUCGAGCUC
CUGCACGCGCCGGCCACGGUUUGCGGGCCA
~~~

#### Linear Design (CAI ≥ 0.8). CAI: 0.808

~~~
AUGUGGCUGGUGAGCCUGGCCAUUGUGACCGCCUGUGCGGGCGCAACCAACUUGUGCCCGUUCGGGGAGGUCUUC
AAUGCGACCAGGUUCGCCAGCGUGUACGCCUGGAAUCGCAAGCGAAUCAGCAAUUGCGUCGCAGAUUAUUCAGUU
UUGUACAACUCCGCCAGCUUUAGCACUUUCAAGUGCUAUGGCGUGAGUCCUACAAAGCUGAAUGAUCUGUGCUUC
ACCAACGUAUAUGCUGAUUCGUUUGUGAUUCGGGGCGACGAGGUCCGGCAGAUCGCGCCGGGGCAGACGGGGAAG
AUAGCAGAUUAUAACUACAAGCUGCCUGAUGAUUUCACUGGAUGUGUCAUCGCUUGGAAUUCGAACAACCUGGAC
AGCAAGGUGGGAGGUAAUUACAAUUACCUCUACCGGCUGUUCAGGAAGUCGAAUCUGAAGCCCUUCGAGCGAGAC
AUAUCCACUGAGAUCUAUCAGGCAGGCUCCACCCCGUGCAACGGGGUGGAGGGUUUUAACUGCUAUUUUCCCCUG
CAGAGCUACGGAUUUCAGCCCACCAACGGGGUGGGGUACCAGCCCUACCGCGUGGUGGUGCUGAGUUUCGAGCUG
CUGCAUGCCCCGGCGACGGUCUGCGGACCU
~~~

#### Our Method (CAI ≥ 0.8). CAI: 0.801

~~~
AUGUGGCUGGUGAGCCUGGCCAUUGUGACCGCCUGUGCGGGCGCAACCAACUUGUGCCCGUUCGGGGAGGUCUUC
AAUGCGACCAGGUUCGCCAGCGUGUACGCCUGGAAUCGCAAGCGAAUCAGCAAUUGCGUCGCAGAUUAUUCAGUU
UUGUACAACUCCGCCAGCUUUAGCACUUUCAAGUGCUAUGGCGUGAGUCCUACAAAGCUGAAUGAUCUGUGCUUC
ACCAACGUAUAUGCUGAUUCGUUUGUGAUUCGGGGCGACGAGGUCCGGCAGAUCGCGCCGGGGCAGACGGGGAAG
AUAGCAGAUUAUAACUACAAGCUGCCUGAUGAUUUCACUGGAUGUGUCAUCGCUUGGAAUUCGAACAACCUGGAC
AGCAAGGUGGGAGGUAAUUACAAUUACCUCUACCGGCUGUUCAGGAAGUCGAAUUUGAAGCCUUUCGAGCGAGAC
AUAUCCACUGAGAUCUAUCAGGCAGGCUCCACCCCGUGCAACGGGGUGGAGGGUUUUAACUGCUAUUUUCCCCUG
CAGAGCUACGGAUUUCAGCCCACCAACGGGGUGGGGUACCAGCCCUACCGCGUGGUGGUGCUGAGUUUCGAGCUC
CUGCAUGCCCCGGCGACGGUCUGCGGACCU
~~~

### EGFP

#### Amino Acid Sequence

~~~
MVSKGEELFTGVVPILVELDGDVNGHKFSVSGEGEGDATYGKLTLKFICTTGKLPVPWPTLVTTLTYGVQCFSRY
PDHMKQHDFFKSAMPEGYVQERTIFFKDDGNYKTRAEVKFEGDTLVNRIELKGIDFKEDGNILGHKLEYNYNSHN
VYIMADKQKNGIKVNFKIRHNIEDGSVQLADHYQQNTPIGDGPVLLPDNHYLSTQSALSKDPNEKRDHMVLLEFV
TAAGITLGMDELYKRSRDISHGFPPAVAAQDDGTLPMSCAQESGMDRHPAACASARINV
~~~

#### Linear Design (unrestricted). CAI: 0.779

~~~
AUGGUCUCCAAAGGUGAGGAGUUGUUCACCGGGGUGGUGCCAAUACUGGUAGAGCUGGAUGGUGAUGUGAACGGG
CAUAAGUUUUCCGUGAGUGGCGAGGGCGAGGGCGACGCCACUUACGGAAAGCUUACGCUCAAGUUCAUAUGCACC
ACCGGCAAGCUGCCAGUACCUUGGCCCACCCUGGUGACAACCCUCACCUAUGGAGUCCAGUGCUUCAGCAGGUAC
CCCGACCACAUGAAGCAGCACGACUUUUUUAAGUCGGCCAUGCCGGAGGGGUACGUGCAGGAGCGCACCAUCUUC
UUCAAGGAUGAUGGUAACUACAAGACGCGGGCCGAGGUGAAGUUCGAGGGGGACACCUUGGUCAACCGGAUUGAG
UUGAAAGGUAUUGAUUUCAAGGAGGACGGGAACAUCCUCGGUCACAAGCUGGAGUAUAAUUACAACAGCCACAAC
GUCUAUAUUAUGGCGGAUAAGCAGAAGAAUGGCAUCAAAGUCAACUUCAAGAUCCGCCAUAAUAUAGAGGAUGGG
AGUGUUCAGCUGGCAGACCACUACCAGCAGAACACUCCGAUCGGGGAUGGGCCCGUCCUCCUGCCUGACAAUCAU
UACCUUUCAACUCAGUCCGCGUUGUCCAAGGACCCCAACGAGAAGCGGGACCACAUGGUCCUGCUUGAGUUCGUG
ACCGCCGCGGGAAUCACGUUGGGGAUGGACGAGCUCUACAAGAGGAGCCGGGACAUCUCCCACGGAUUCCCGCCG
GCGGUCGCGGCGCAGGAUGACGGGACAUUGCCGAUGUCCUGUGCUCAGGAGUCGGGGAUGGACCGUCAUCCUGCG
GCUUGCGCCUCGGCCCGCAUCAAUGUA
~~~

#### Our Method (unrestricted). CAI: 0.774

~~~
AUGGUCUCCAAAGGUGAGGAGUUGUUCACCGGGGUGGUGCCAAUACUGGUAGAGCUGGAUGGUGAUGUGAACGGG
CAUAAGUUUUCCGUGAGUGGCGAGGGCGAGGGCGACGCCACUUACGGAAAGCUUACGCUCAAGUUCAUAUGCACC
ACCGGCAAGCUGCCAGUACCUUGGCCCACCCUGGUGACAACCCUCACCUAUGGAGUCCAGUGCUUCAGCAGGUAC
CCCGACCACAUGAAGCAGCACGACUUUUUUAAGUCGGCCAUGCCGGAGGGGUACGUGCAGGAGCGCACCAUCUUC
UUCAAGGAUGAUGGUAACUACAAGACGCGGGCCGAGGUGAAGUUCGAGGGGGAUACCUUGGUCAACCGGAUUGAG
UUGAAAGGUAUUGAUUUCAAGGAGGACGGGAACAUCCUCGGUCACAAGCUGGAGUAUAAUUACAACAGCCACAAC
GUCUAUAUUAUGGCGGAUAAGCAGAAGAAUGGCAUCAAAGUCAACUUCAAGAUCCGCCAUAAUAUAGAGGAUGGG
AGUGUUCAGCUGGCAGACCACUACCAGCAGAACACUCCCAUCGGGGAUGGGCCCGUCCUCCUUCCUGAUAAUCAU
UACCUUUCAACUCAGUCCGCGUUGUCCAAGGACCCCAACGAAAAGCGGGACCACAUGGUCCUGCUUGAGUUCGUG
ACCGCGGCGGGAAUCACGUUGGGGAUGGACGAGCUCUACAAGAGGAGCCGGGACAUCUCCCACGGAUUCCCGCCG
GCGGUCGCGGCGCAGGAUGACGGGACAUUGCCGAUGUCCUGUGCUCAGGAGUCGGGGAUGGACCGUCAUCCUGCG
GCUUGCGCCUCGGCCCGCAUCAAUGUA
~~~

#### Linear Design (CAI ≥ 0.8). CAI: 0.806

~~~
AUGGUCUCCAAGGGUGAGGAGUUGUUCACCGGGGUGGUGCCAAUACUGGUAGAGCUGGAUGGUGAUGUGAACGGG
CAUAAGUUUUCCGUGAGUGGCGAGGGCGAGGGCGACGCCACUUACGGAAAGCUUACGCUCAAGUUCAUAUGCACC
ACCGGCAAGCUGCCAGUACCUUGGCCCACCCUGGUGACAACCCUCACCUAUGGAGUCCAGUGCUUCAGCAGGUAC
CCCGACCACAUGAAGCAGCACGACUUUUUUAAGUCGGCCAUGCCGGAGGGGUACGUGCAGGAGCGCACCAUCUUC
UUCAAGGAUGAUGGUAACUACAAGACCCGGGCGGAGGUGAAGUUCGAGGGGGACACCUUGGUCAACCGGAUUGAG
UUGAAAGGUAUUGAUUUCAAGGAGGACGGGAACAUCCUCGGCCACAAGCUGGAGUAUAAUUACAACAGCCACAAC
GUCUAUAUUAUGGCGGAUAAGCAGAAGAAUGGCAUCAAAGUCAACUUCAAGAUCCGCCAUAAUAUAGAGGAUGGG
AGUGUUCAGCUGGCAGACCACUACCAGCAGAACACUCCCAUCGGGGAUGGGCCCGUCCUCCUGCCUGACAAUCAU
UACCUUUCAACUCAGUCCGCGUUGUCCAAGGACCCCAACGAAAAGCGGGACCACAUGGUCCUGCUUGAGUUCGUG
ACGGCCGCGGGGAUAACCCUGGGGAUGGACGAGCUCUACAAGAGGAGCCGGGACAUCUCCCACGGGUUUCCCCCG
GCCGUCGCGGCGCAGGACGAUGGGACCCUUCCCAUGUCCUGCGCGCAGGAGAGCGGGAUGGACCGCCAUCCCGCU
GCCUGCGCCUCCGCCCGGAUCAAUGUA
~~~

#### Our Method (CAI ≥ 0.8). CAI: 0.800

~~~
AUGGUCUCCAAGGGUGAGGAGUUGUUCACCGGGGUGGUGCCAAUACUGGUAGAGCUGGAUGGUGAUGUGAACGGG
CAUAAGUUUUCCGUGAGUGGCGAGGGCGAGGGCGACGCCACUUACGGAAAGCUUACGCUCAAGUUCAUAUGCACC
ACCGGCAAGCUGCCAGUACCUUGGCCCACCCUGGUGACAACCCUCACCUAUGGAGUCCAGUGCUUCAGCAGGUAC
CCCGACCACAUGAAGCAGCACGACUUUUUUAAGUCGGCCAUGCCGGAGGGGUACGUGCAGGAGCGCACCAUCUUC
UUCAAGGAUGAUGGUAACUACAAGACCCGGGCGGAGGUGAAGUUCGAGGGGGAUACCUUGGUCAACCGGAUUGAG
UUGAAAGGUAUUGAUUUCAAGGAGGACGGGAACAUCCUCGGUCACAAGCUGGAGUAUAAUUACAACAGCCACAAC
GUCUAUAUUAUGGCGGAUAAGCAGAAGAAUGGCAUCAAAGUCAACUUCAAGAUCCGCCAUAAUAUAGAGGAUGGG
AGUGUUCAGCUGGCAGACCACUACCAGCAGAACACUCCCAUCGGGGAUGGGCCCGUCCUCCUUCCUGAUAAUCAU
UACCUUUCAACUCAGUCCGCGUUGUCCAAGGACCCCAACGAAAAGCGGGACCACAUGGUCCUGCUUGAGUUCGUG
ACGGCCGCGGGGAUAACCCUGGGGAUGGACGAGCUCUACAAGAGGAGCCGGGACAUCUCCCACGGGUUUCCCCCG
GCCGUCGCGGCGCAGGACGAUGGGACCCUUCCCAUGUCCUGCGCGCAGGAGAGCGGGAUGGACCGCCAUCCCGCU
GCCUGCGCCUCCGCCCGGAUCAAUGUA
~~~

